# Label-free single-instance protein detection in vitrified cells

**DOI:** 10.1101/2020.04.22.053868

**Authors:** J. Peter Rickgauer, Heejun Choi, Jennifer Lippincott-Schwartz, Winfried Denk

**Affiliations:** Howard Hughes Medical Institute, Janelia Research Campus, Ashburn, VA, USA; Max-Planck Institute of Neurobiology, Martinsried, Germany

## Abstract

A general method to map molecular interactions and conformational states in structurally intact cells would find wide application in biochemistry and cell biology. We used a library of images— calculated on the basis of known structural data—as search templates to detect targets as small as the “head” domain (350 kDa) of the ribosome’s small subunit in single-tilt electron cryo-micrographs by cellular high resolution template matching (cHRTM). Atomically precise position and orientation estimates reveal the conformation of individual ribosomes and enable the detection of specifically bound ligands down to 24 kDa. We show that highly head-swivelled states are likely to play a role in mRNA translocation in living cells. cHRTM outperforms cryo-electron tomography three-fold in sensitivity and completely avoids the vicissitudes of exogenous labelling.

---

Finding molecular components in intact cells has driven the development of biological imaging techniques, some of which use specific labels, such as fluorescent antibodies, while others rely on a target’s intrinsic properties such as its shape (Oliver 1973) and structure (Rickgauer, Grigorieff, and Denk 2017). Labelling is frequently incomplete or nonspecific, can alter the target’s interactions with other molecules, provides only moderate precision of localization and orientation, and can simultaneously track only a limited number of targets. Label-free cryo-electron tomography (CET) is routinely used to analyze macromolecular complexes (Asano, Engel, and Baumeister 2016) and can identify some of their building blocks based on their intrinsic shape, but struggles with components below a molecular weight (MW) of about one megadalton (MDa) because very fine structural features are destroyed by radiation damage accumulated over a tilt-series (Oikonomou, Chang, and Jensen 2016). Proteins contain structural features down to well below one nanometer, and for many proteins, maps of the electron density or lists of atomic coordinates are available at a resolution of 0.3 nm or better. As long as motion blurring is avoided by recording images at a sufficiently high frame rate followed by alignment (Brilot et al. 2012), sample structure at that resolution can be propagated with little attenuation in the transmission electron microscope (TEM), and has been shown to be essential for the sensitive and specific detection of proteins by high-resolution template matching (HRTM) (Rickgauer, Grigorieff, and Denk 2017).

HRTM uses a library of templates, each sensitive to the target at a particular orientation and sample depth. Every peak detected in the cross correlogram (CCG) comes with location and orientation estimates, far exceeding in precision (see below) that of any antibody or fluorescence-based technique. Here we introduce cellular HRTM (cHRTM), the extension of HRTM to vitrified cells, and show that it can: a) detect individual ribosomes and determine their conformation up to a sample thickness of several hundred nanometers, b) identify intermolecular interactions by finding one protein frequently in the same orientation with respect to another, and c) detect specifically bound ligands as small as 24 kDa by restricting the search space on the basis of prior information about a target’s possible location and orientation.

We imaged structurally intact vitrified mouse embryonic fibroblast (MEF) cells using cryo-electron microscopy (cryo-EM), accepting only zero-loss electrons for detection to eliminate shot noise due to inelastic scattering (Fig. 1A,B). To prevent a loss of resolution and, more importantly, a loss in HRTM-sensitivity by motion blurring, image data were acquired as a sequence of frames in quick succession (a dose-fractionated movie), later realigned, combined into a single image (Brilot et al. 2012), and searched (Fig. 1C).

**Fig. 1.**
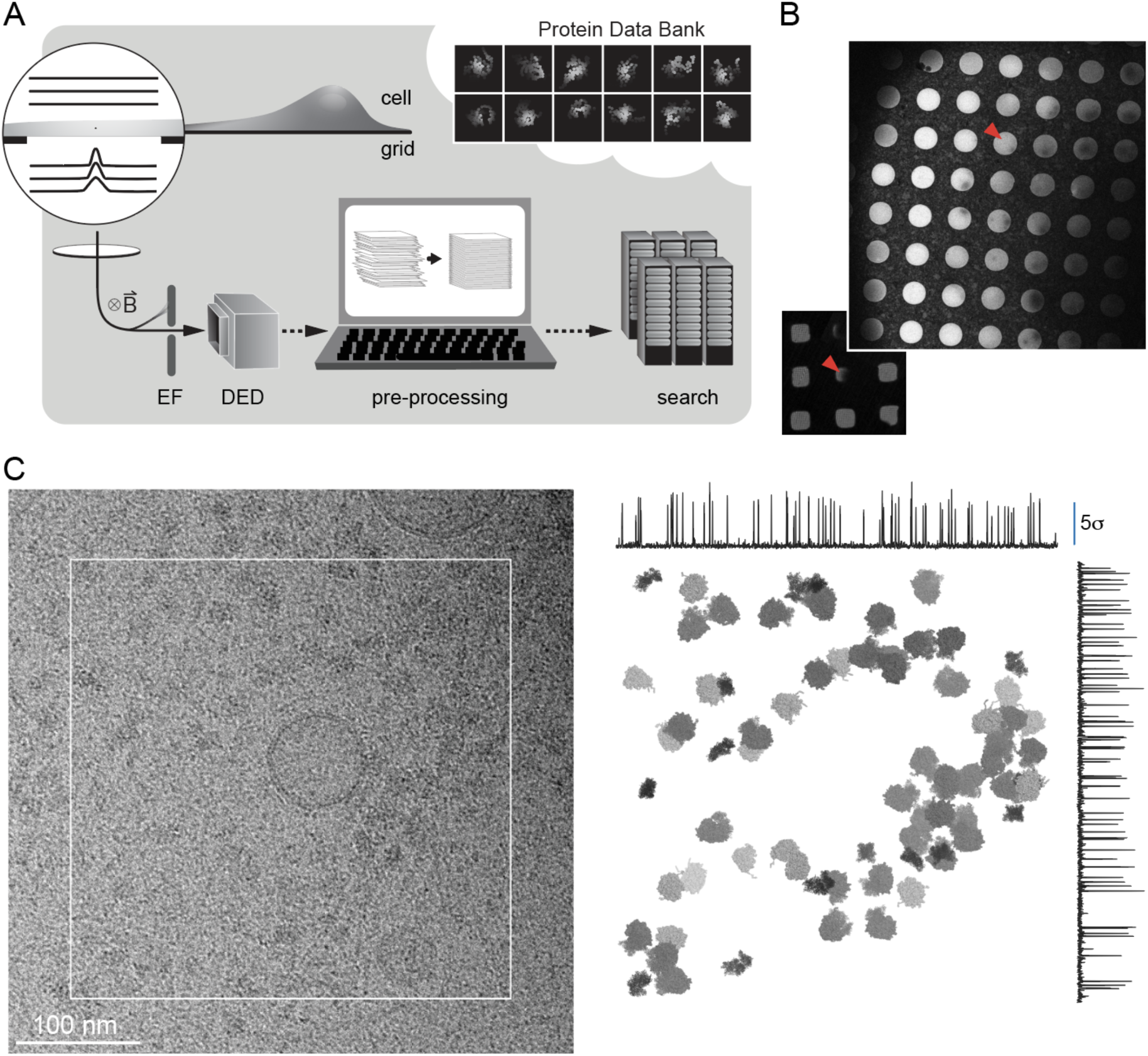
Data acquisition and analysis. **A)** cHRTM workflow. Counterclockwise, starting from upper left: Cell growing on EM sample grid; zoomed: electron wave passing grid opening; electron optics; energy filter; direct electron detector; data-preprocessing; search, using known structural data. **B)** Cells grown on EM grids and imaged in the TEM after plunge-freezing. Low magnification (behind; grid pitch is 85 μm), showing grid openings (red arrowheads indicate centers of zoomed images), and at medium magnification, showing openings in the gold film (hole period 2.5 μm). **C)** TEM image at 3 μm underfocus (left); white square: FOV of the image taken earlier for cHRTM. Right: targets found by cHRTM and represented by to-scale models at the correct locations and orientations. Dark, medium, and light gray: SSU, SSU+LSU, and LSU particles. Line plots of cross correlations, maximum-projected across orientations, and either the horizontal or vertical spatial direction.

Central to HRTM is the creation of a library of high-resolution templates covering all possible target orientations and depths. Each template is a noise-free but otherwise realistic image of the target, calculated using the target’s known atomic structure and the optical properties of the microscope. The number of templates ranged from one, for ligand detection, to tens of millions, when searching a thick sample without any orientation restrictions. The number of cross-correlation values produced in the latter case can approach 10^15^, but after preselection by SNR and clustering by spatial and orientational proximity (to avoid instance duplication), the number of peak candidates remaining was small enough to permit each peak’s position, depth, orientation and SNR value estimates to be to be optimized. This reduced sampling-phase dependent peak-value fluctuations (Fig. 2A). We applied a flat-fielding technique (adapted from Ref. (Afanasyev et al. 2015)) to compensate for spatial inhomogeneities in the CCG, caused by variations in the macromolecular density across the FOV, which, as searches for a decoy target show, restored Gaussian noise behaviour (Fig. 2B, Supplemental Fig. 1, see Methods).

**Fig. 2.**
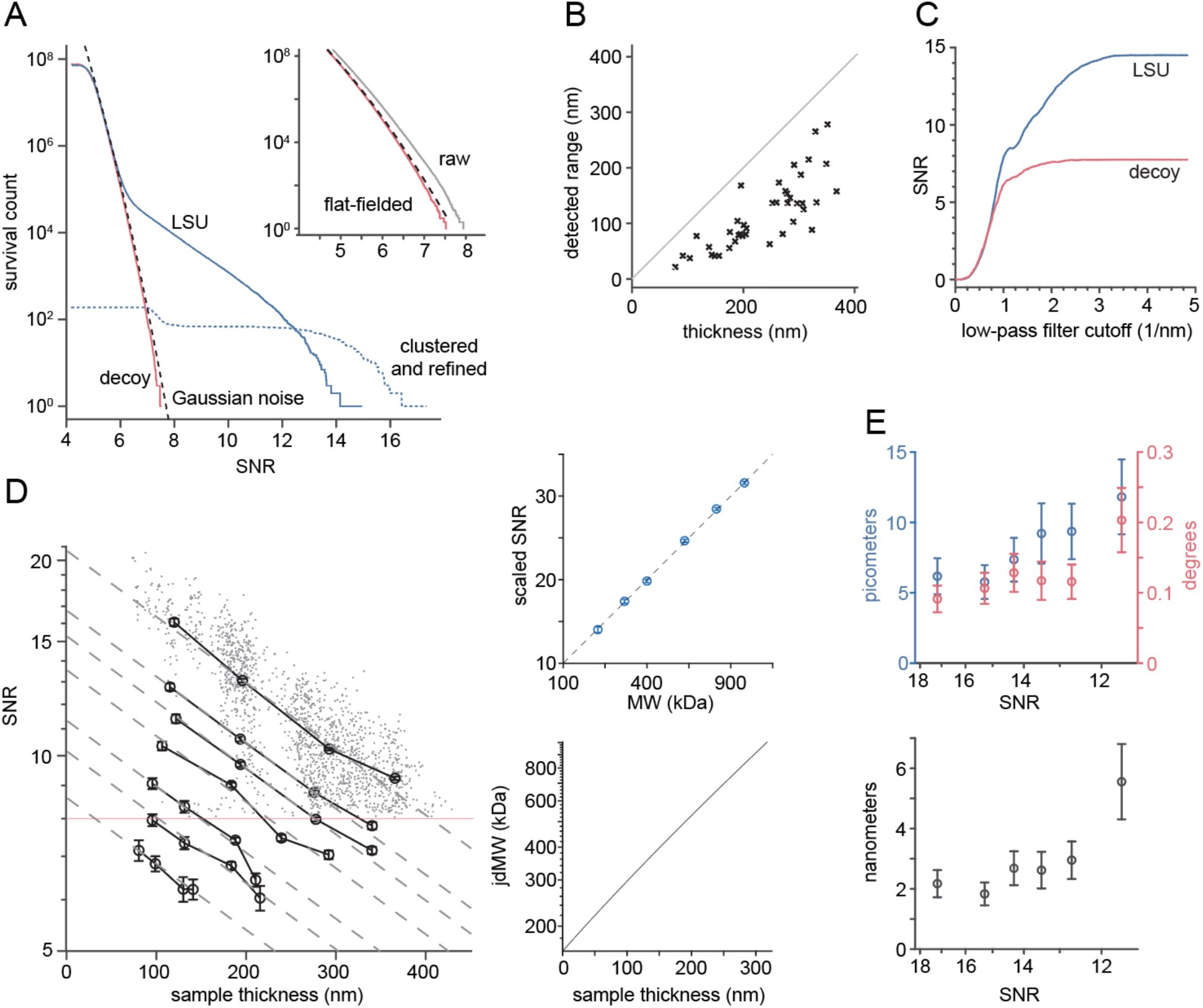
Detection sensitivity and precision. **A)** Distribution of CC values (for one FOV) as a survival histogram for decoy (red), LSU before (blue), and after (dotted) clustering and refinement. Inset: Distributions of CC values for a decoy search before (gray) and after (red) flat-fielding; Gaussian noise model (dashed). **B)** Depth range of detected LSUs *vs*. sample thickness. **C)** LSU-SNR *vs*. spatial low-frequency filter roll-off (blue; average of 10 peaks) and decoy (red). **D)** Left: Average SNR for 4 sample thickness bands, which together span the thickness range, *vs*. sample thickness (means across each band) for centered spherical LSU fragments (200, 300, 400, 600, 800, and 1000 kDa in MW, 4.34 to 7.84 nm in radius) and the entire LSU (top trace); dashed lines: exponential decay curves (all with a λ_SNR_=426 nm decay constant) fit to fragment data. Thin red line indicates a typical detection threshold used for unconstrained searches (fragment SNR values were determined at the location and orientation found for the LSU from which they were derived). Top right: SNR (scaled to compensate for thickness) *vs*. fragment MW (open circles, showing mean and SEM; dashed line, SNR=sqrt(MW)). Lower right: just detectable molecular weight (kDa) *vs*. sample thickness. **E)** Upper: lateral and orientational uncertainties measured using the de-interlaced-movie method *vs*. SNR, with 1/SNR increasing linearly along the x-axis; lower: optimized-focus uncertainty.

Searching 547 images of the cellular periphery with the ribosomal large subunit (LSU) as the target, we found a total of 9,359 peaks, many extending well above the noise floor (Fig. 2A). LSUs were found in sample locations as thick as ∼400 nm and were distributed across most of the sample depth (Fig. 2B). Since each template is only effective (Rickgauer, Grigorieff, and Denk 2017) over a depth of about 40 nm, an exhaustive search of a 200 nm thick sample required 5 different templates for each orientation (Fig. 2B). The dependence of the SNR on the range of spatial frequencies included shows that cHRTM needs the fine details of the target’s atomic structure to reliably detect instances embedded in the crowded molecular environment of a cell (Fig. 2C).

As the sample thickness increases, both inelastic and wide-angle scattering deplete the set of coherent electrons contributing to phase contrast, on which cHRTM depends. This reduction limits the achievable SNR, which scales with the square root of the electron number and should thus decay with thickness at least as fast as an exponential with twice the coherent electrons’ decay length (2 × 283 nm; Supp. Fig. 2). Using as search targets artificial spherical fragments of the LSU, we experimentally determined the dependence of the SNR on thickness (T) and MW and found that it closely follows the expression 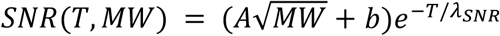, with a SNR decay length, λ_SNR_, of 426 nm (Fig. 2D). SNR loss beyond that expected from inelastic and wide-angle scattering (Methods) is likely due to structural noise caused by macromolecular background. The resulting just detectable MW (jdMW) approximately doubles with every increase in thickness by 125 nm (Fig. 2D). At 100 nm the jdMW is 296 kDa, matching the prediction (Rickgauer, Grigorieff, and Denk 2017) that in a 100 nm thick region of a cell it should be possible to detect targets with a MW above 300 kDa.

To determine location and orientation precision for cHRTM we de-interlaced single movies into two datasets, which contain the same targets but uncorrelated noise, and searched them independently for LSUs. Location and orientation discrepancies within pairs of peaks corresponding to the same target were then used to estimate localization and orientation errors, which when extrapolated to the undivided dataset yield (for a SNR of 14) 8.0 pm and 0.125°, respectively (Fig. 2E). Note that this estimate applies only to combinations of SNR and search space where outliers are rare enough to be ignored.

Vitrification arrests macromolecular dynamics “midflight” (Moore 2012), creating a preparation that contains all reaction intermediates in their states at the time of freezing. For reactions that are ongoing and unsynchronized, conformations are sampled without bias, i.e., every state’s prevalence is proportional to the average time spent in that state. As a first application, we considered the process of polypeptide chain elongation, a key step in protein synthesis and the central activity of ribosomes in cells. Evidence *in vitro* suggests (for reviews see refs. (Larsen et al. 2019; Ramakrishnan 2014)) that elongation requires precise relative movements among the three parts of the ribosome (LSU, head, and body, Fig. 3A), presumably to ensure that the space inside the ribosomal cavity has the right shape for the reactants to assemble in a conformation suitable for the current reaction step, while still providing the confinement needed to suppress side reactions that might lead to faulty proteins. However, structural data in support of a mechanistic description of ribosome function have been obtained almost entirely using ribosomes extracted from cells, often trapped by chemical manipulation in a particular state (Frank 2017). Such preparations yield high-resolution structural data but do not tell us how often, if ever, a ribosome assumes these configurations in living cells.

**Fig. 3.**
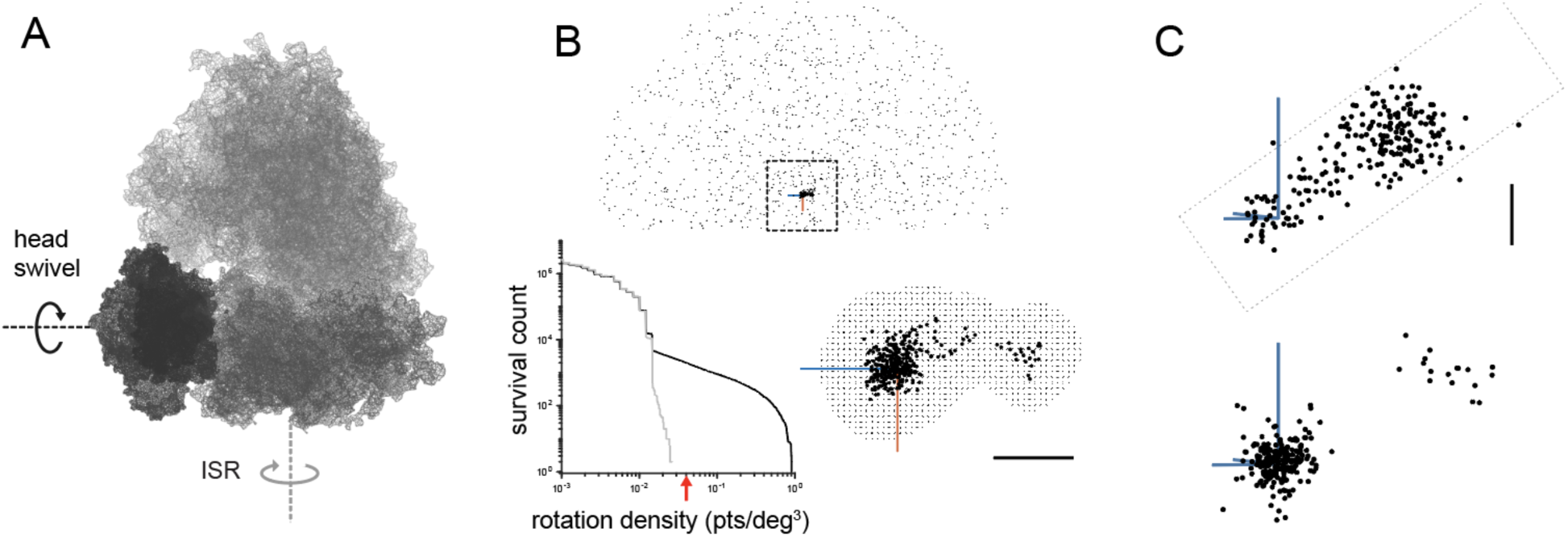
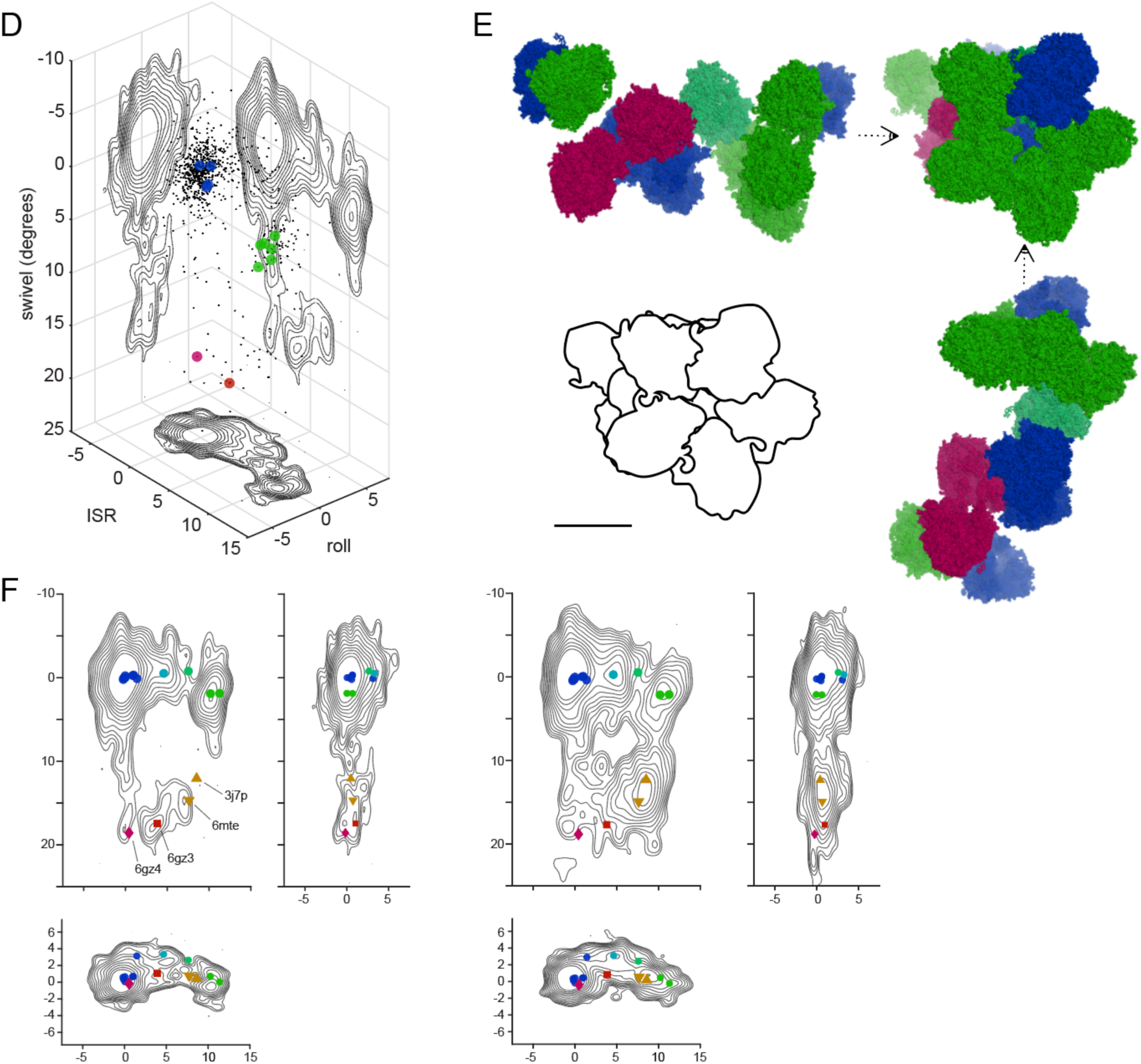
Range of ribosome conformations. **A**) LSU (top); body (lower right) and head (lower left). Axes for intersubunit rotation (ISR) and head swivel as indicated. **B**) Top: Relative orientations (ROs) of LSU and body for individual instances represented as rotation vectors. Lower left: distribution (as survival histograms) of the LSU/body relative-orientation density (black) compared to that for the case without any correlation between orientations as control (gray). Red arrow: density threshold used. Lower right: RO density thresholded points and the resulting search mask after padding by 2°. **C**) ROs between head and body without (top) and with (bottom) an LSU bound. **D**) Conformations for 847 LSU instances (black dots) with smoothed kernel densities projected onto the walls and contour plotted. Transparent spheres: polysomal ribosomes using the same colors as in (E). **E)** Orthogonal views of the same polysomal array. Models placed using cHRTM derived positions (x,y, and LSU depth) and orientations (Methods). **F)** Ribosome conformation densities for normal (left) and starved (right) cells, projected onto the rotation *vs*. swivel plane (top), the roll *vs*. rotation plane (bottom) and the swivel *vs*. roll plane (right). Contour lines are spaced by a constant factor of sqrt(2) with the lowest line corresponding to a density of 0.3 instances/degree^2^. Scale bars are: 5 degrees in (B) and (C), and 25 nm in (E).

To determine the range of native ribosome conformations, we separated a single, highly resolved structural model (Natchiar et al. 2017) into LSU, body, and head. We used a first set of searches, performed without restraints on positions and orientations, to create unbiased maps of preferred relative orientations (ROs) within LSU/body and body/head pairs. For both pairings the ROs were concentrated in clearly delineated regions, easily seen when displaying ROs as rotation vectors (Fig. 3B). The peaks of the density corresponded closely to the (unrotated) conformation of the reference structure (Fig. 3B,C), and the surrounding densities, presumably, belong to conformations assumed during various stages of the elongation cycle. Using RO density thresholds to restrict searches for bodies and then for heads, using the bodies found, enabled us to reliably detect heads in locations as thick as 370 nm and resulted in a dataset with 226 free SSUs (head and body detected, but no nearby LSU) and 847 fully assembled ribosomes.

It quickly became clear as we started to analyze this dataset that, across all SSUs, irrespective of their relationship with any LSU, the head/body ROs were mostly confined to a cylindrical volume (radius= 5°), spread out along the cylinder’s axis over a length of about 21° (Fig. 3C). The cylinder’s axis was in close agreement (3.1° deviation) with the swivel axis reported for eukaryotic ribosomes *in vitro* (Flis et al. 2018). Free SSUs varied roughly uniformly in the rotation between body and head, while LSU-bound SSUs were polarized into canonical (un-swiveled; n=202) and swiveled states (n=17), just over 16° apart (Fig. 3C). These observations suggest that the total swivel range is limited by steric interactions within the SSU, but that interactions of the head with other parts of the assembled ribosome control the position inside that range (Gabashvili et al. 1999).

The conformations determined for a set of complete ribosomes shows the extent of structural variation that exists in living cells (Fig. 3D; 1731 LSUs, 1134 with bodies, 99% precision; 847 with heads, 95% precision). Each conformation can be mapped back to the ribosome’s precise location to, for example, test if a ribosome’s membership in a complex super-assembly such as a polysome (Fig. 3E) affects conformational prevalence. While polysomes have been studied extensively with electron microscopy, initially with negative stain (Warner, Knopf, and Rich 1963), even recent studies with CET (Brandt et al. 2010; Afonina and Shirokov 2018) have struggled to detect single instances of the small subunit reliably, let alone the head, leaving one of the ribosome’s main conformational parameters inaccessible.

To test and refine the map between the elongation cycle’s reaction coordinate and the space of conformations, we determined where on the ribosome and for which conformations some of the essential reactants and cofactors of the elongation reaction are bound. The ligands we explored range in size from 24-80 kDa (tRNA; elongation factor 2, EF2), well short of the MW required for reliable detection without prior information (Fig. 2D). Combining high-resolution structures of ligands bound to particular sites on the ribosome (Methods) with the highly precise locations and orientations from cHRTM provides the information necessary to constrain the search space sufficiently to detect single instances in favorable cases (Fig. 4A). Because single-instance ligand detection was possible only for ribosomes in very thin sample regions, a small fraction of our dataset, we grouped ribosomes by conformation and averaged the ligand signals, which allowed us to test a wider collection of ligands including multiple structures of tRNA in different conformations and binding sites (Fig. 4, Supp. Fig. 4).

**Fig. 4.**
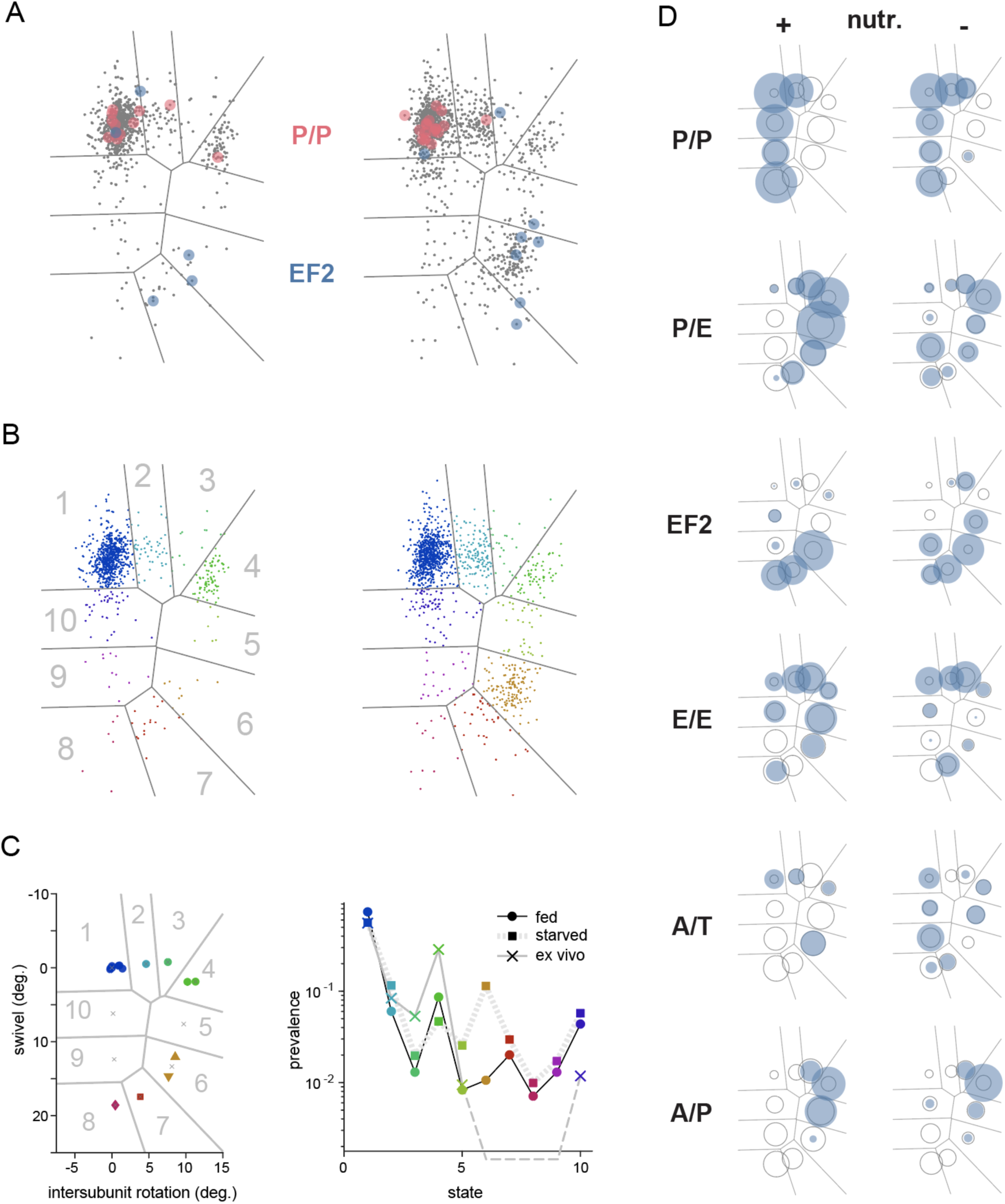
Ligand binding. **A)** Single-instance ligand detection of P/P-site tRNA and EF2. **B)** Ribosome instances grouped by proximity to known conformations for fed (left) and starved (right) cells. Gray lines: region boundaries. **C)** Left: region key. Right: Prevalence of ribosome conformations in fed and starved cells *vs*. region index. Also shown are prevalences obtained from a simulation of Ref. (Behrmann et al. 2015) data (Suppl. Fig. 5). **D)** Ribosome-bound tRNA in the P/P, P/E, E/E, A/T, and A/P state as well as elongation factor 2 for fed (left) and starved (right) cells. Solid disks: Estimated occupancy proportional to disk area. The area of open circles indicates the SD. The highest occupancy (0.9) was for P/E in area 5 (fed conditions).

Most ribosomes (∼74%) were found in the unrotated state (near the origin; region 1 in Fig. 4B), a conformation in which bound tRNA was also detected in classical (P/P and E/E) and codon-recognition (A/T) configurations. As expected for actively translating ribosomes, the fractional binding-site occupancy in this region is high for P/P-site tRNA (0.6; Fig. 4D). Because target distortion and misalignment reduce the estimated occupancy below the actual value, this result may still be consistent with the notion that the LSU’s P-site is rarely empty (Behrmann et al. 2015). (Note that we use *prevalence* for the distribution of ribosomes among conformations but *occupancy* for the fraction of binding sites with a ligand.) The picture is substantially different for ribosomes in severely nutrient-deprived (starved) cells, where region-1 prevalence is only about 56%. Ribosomes in a state with high ISR, low head swivel (<5°), and tRNA bound in hybrid configurations (A/P and P/E; region 4) are half as prevalent as in normal cells (5% vs. 9%), a ratio that is reversed (12% vs. 6%) in region 2, the “rolled” conformation (Budkevich et al. 2014).

Conformations with high head-swivel angles (regions 6-9) together contain only 5% of all instances under normal conditions but 17% under starved conditions, and are the only regions with significant binding of EF2. Detection of highly head-swiveled instances with bound EF2, a protein necessary for a eukaryotic ribosome’s advance to the next mRNA codon (Flis et al. 2018), demonstrates that cHRTM can capture short-lived reaction intermediates, apparently lost outside the cell (Behrmann et al. 2015). While the function of a ribosome’s head swivel remains a matter of debate, structural snapshots of stalled ribosomes at high head-swivel (Agrawal et al. 1999; Frank and Agrawal 2000; Flis et al. 2018) have suggested that such a state could allow the movement of the mRNA-tRNA complex during translocation (Ratje et al. 2010). Our observation that a significant fraction of ribosomes in intact mammalian cells are in a head-swiveled state supports the idea that this transient conformation is an essential intermediate during translation.

It came as a surprise that cell growth under starvation substantially shifted the highest ISR ribosomes from an un-swivelled to a highly swivelled state, gaining EF2 in the process of arriving at conformations seen for inactive (empty) ribosomes (Brown et al. 2018), which suggests that EF2, which is essential for productive translation in eukaryotic cells (Flis et al. 2018), may in addition suppress translation under some conditions (Brown et al. 2018).

Given a cycle time of ∼200 ms for elongation (Yan et al. 2016), a delay of seconds (let alone minutes) would certainly change prevalences of functional states, as reactions continue until they encounter a step requiring one of the reactants lost upon cell lysis. This may explain why the highly-swivelled head conformations escaped detection in a study specifically designed to capture a complete set of states visited during the elongation cycle (Behrmann et al. 2015). Even from a small number of instances, as long as they are detected reliably, one can infer with high confidence the existence of elusive and unstable states and estimate their prevalence. Because cHRTM requires only single images, it should be straightforward to acquire datasets with hundreds of thousands of instances and to screen them with high detection precision for small groups with distinct but highly transient conformations and lifetimes in the microsecond range.

The need for only a single un-tilted image greatly simplifies acquisition and makes cHRTM compatible with pulsed-beam (Spence 2017) and multi-pass imaging techniques (Juffmann et al. 2017; Koppell et al. 2019), both designed to increase the amount of high-resolution information extracted before it is lost to radiation damage. Together with improved detectors, this might well boost the sensitivity of cHRTM enough to make the majority of proteins detectable. The jdMW is inversely proportional to the allowable electron exposure and currently stands at around 300 kDa for unconstrained cHRTM searches (Fig. 2D), about a factor of three below what we have seen reported for CET, which is fundamentally incompatible with single-shot imaging and may not be compatible with multi-pass imaging, due to CET’s reliance on multiple images at changing sample tilt. We expect that a carefully designed hybrid approach will eventually combine the strengths of 2D cHRTM and CET, with the first image taken at zero tilt and used for cHRTM.

## Materials and Methods

### Cell culture and sample preparation

Immortalized mouse embryonic fibroblast (MEF) cells (Lionnet et al. 2011) were cultured in DMEM supplemented with 10% (v/v) inactivated FBS (Corning), 2 mM glutamine, penicillin (100 IU/mL), and streptomycin (100 µg/mL) at 37 °C and 5% CO2. Cells were starved by switching the culture medium to HEPES-buffered Hank’s Balanced Salt Solution (HBSS) 10 hours before vitrification. For TEM imaging, cells were plated and cultured for 24 h on gold-mesh grids before freezing. Grids were coated with either holey gold foil (UltrAuFoil R1.2/1.3 or R0.6/1.0, Electron Microscopy Sciences) or a continuous ultrathin carbon film on a lacey carbon support (#01824G, Ted Pella). To remove any residual organic contamination and render the grids hydrophilic, they were first exposed to a low pressure oxygen plasma for 30 sec at −15 mA using an Easiglow plasma cleaner (Ted Pella), then immersed in a solution of Human Plasma Fibronectin (10 ug/mL in PBS; #FC010, EMD Millipore), and washed thoroughly in PBS. After placing grids with their foiled sides up in a custom teflon grid carrier, which was attached with Kwik-cast (WPI) to the bottom of a 35 mm cell culture dish, and immersing them in 3 mL supplemented DMEM, 1 mL of cells in suspension (100,000-200,000 cells) was added dropwise to the dish. The dish was transferred to an incubator (37 °C, 5% CO2) and after 24 h topped up with pre-warmed supplemented DMEM and carefully moved to a second incubator located near the plunge-freezing instrument (Leica EM GP2). The dish was removed from the incubator about 10 min before the start of the plunge freezing procedure, which was completed for each grid before proceeding to another grid and involved 1) immersing the grid for 5 s in phosphate-buffered saline (PBS), 2) blotting excess fluid from the backside of the grid (5 s in a controlled-environment chamber; 37 C, 95% relative humidity), and 3) plunging the grid into liquid ethane (−173 C). After vitrification, grids were rapidly transferred into liquid nitrogen, where they were stored until imaging.

### Data acquisition

All images were acquired at 300 kV electron energy using a TEM (Titan Krios, FEI/Thermo) equipped with a cryogenic stage and automated sample changer, a corrector for spherical aberration, a post-column energy filter (Gatan, set to admit electrons with a loss of up to 10 eV), and direct electron detector (Gatan, model K2). Data for cHRTM were recorded as dose-fractionated movies in super-resolution mode at 20 Hz frame rate and at a nominal pixel pitch of 0.1048 nm, using exposure rates between 600 and 800 e/(nm^2^s), parallel illumination (condenser aperture C2 diameter = 50 *μ*m), and with the 100 *μ*m diameter objective aperture, which eliminates spatial frequencies >10 cycle/nm. Low-dose techniques were used throughout, in particular when scouting potential target areas.

To find promising locations, we first selected, using low magnification image montages (pixel size 1.98 um), grid openings for which an intensity gradient near the edges indicated the presence of cells, which were then viewed at higher magnification (pixel size 9.53 or 2.76 nm), and locations with an intact support film, limited ice contamination, and a brightness consistent with a sample thickness below ∼500 nm were chosen as target areas. At each target area, the stage height was adjusted so that the rotation axis was close to the sample plane (eucentric condition), refined using eucentric focusing routines (coarse and fine) in SerialEM (Mastronarde 2018), and then mapped using image montages acquired at the same magnification. We stitched these medium-magnification montages (MMMs) together using Blendmont (IMOD) to generate an attenuation-based thickness map, which we then used to select a number of areas with a thickness below ∼500 nm for high-resolution imaging. For each square, ∼10 to 100 image-center locations were chosen. Two movies were acquired at each location: first, a high-resolution movie with a target underfocus of 300 nm, for HRTM, followed by a “context” movie at higher defocus and lower magnification.

### Data pre-processing

Super-resolution movie frames were gain-corrected, purged of “hot” pixels, corrected for anisotropic magnification distortion (using the *mag_distortion_estimate* and *mag_distortion_correct* software (Grant and Grigorieff 2015), with distortion parameters (typically <2%) estimated from images of a polycrystalline gold reference grid, and resampled to a final pixel size of 0.1032 nm. To estimate shift vectors for movie frame alignment, the movie was downsampled to a pixel size of 0.2064 nm before cross-correlating all frame pairs. The center patches (32×32) of all cross correlograms (CCGs) with frame index differences >20 were averaged and subtracted from all the other CCG patches to eliminate fixed pattern noise. An initial set of shift vectors (one for each frame, except the first one) was generated by accumulating incremental shift vectors based on the locations of the maximum values for each of the CCGs between consecutive frames. We then used optimization to find the set of shift vectors that maximizes the central peak in the sum of shifted CCGs between consecutive frames. The obtained set of shift vectors was then applied to the unbinned movie and all frames taken during the first ∼1500 e/nm^2^ of exposure were summed and MTF-corrected, yielding an image ready for template matching (Grant and Grigorieff 2015; Rickgauer, Grigorieff, and Denk 2017). Note that the MTF includes a dose rate-dependent term (Ruskin, Yu, and Grigorieff 2013). CCG-patch and movie-frame shifts were implemented by phase-ramp multiplication in Fourier space to allow sub-pixel displacements.

### Searching

HRTM searches were performed essentially as described (Rickgauer, Grigorieff, and Denk 2017), with the following differences. 1) We performed several searches of each image, which used, in addition to the “average” defocus value as estimated using CTFFind4 (Rohou and Grigorieff 2015), a number of values above and below (typically in 50 nm increments). 2) Offset and gain were flat-fielded (Afanasyev et al. 2015) pixel-wise by first subtracting the mean and then dividing by the standard deviation of all CC values across orientations. 3) CC-cluster peak pixel values were identified by first taking, pixel-wise, the maximal CC value across all templates (orientations and defocal planes). Values above a threshold of 7.0 were clustered using 8 connectivity, and clusters with peak pixels spaced < 10 pixels and < 10° were merged. Using a square patch centered on the peak and large enough to contain the template, we then refined the surviving CC peaks by adjusting for each of them focus and orientation until a local maximum of the CC value was found, by using a sequence of local, refined-grid searches, varying first the focus (+/- 50 nm range, in 4 nm increments), then the orientation (2°, in 0.5° increments), and, again, the focus (−50 to +50 nm, in 2 nm increments). Locations and orientations of peaks were then optimized by simultaneously varying the projected location and orientation using a five-dimensional search (Nelder-Mead simplex algorithm, *fminsearch* function in Matlab R2019a; Mathworks) to determine two phase-ramp parameters (in-plane shifts) and three orientation parameters that maximized the peak amplitude. 4) The HRTM search algorithm was adapted so it could take advantage of GPUs. This reduced the computation time needed ∼2-fold, compared to our earlier implementation on a CPU cluster of similar purchase price. 5) For the calculation of scattering potentials, B-factors for all targets except for the LSU and the decoy were set to zero.

The spherical fragments of the LSU used for evaluating thickness-dependence and the just-detectable molecular weight (jdMW) were isolated from the full model (PDB:6ek0) by selecting atoms based on their Euclidean distance from the LSU’s center of mass, and then expanding the radius of included atoms until the intended MW was reached. The decoy template was generated using a high-resolution model for an *E. Coli* ribosome (PDB:5j5b), and included 147,242 atoms (2,081 kDa). The structure was modified by introducing an offset and rotation between LSU and SSU resulting in an overall shape that when viewed in some projections resembles the true LSU target.

### Searching for complete ribosomal instances

We took the existing searches for LSUs (547 FOVs) and selected, based on the thickness dependence of the SNR, those FOVs for which independent and unrestricted searches for body (690 kDa) and head (350 kDa) were likely to yield (at a threshold of 7) viable candidates in a search for preferred relative orientations (ROs). 87 FOVs met this criterion for bodies, and of those, 34 also for heads. The former were searched for bodies and the latter for heads as well.

Together, the 87 FOVs contained 10,451 LSU/body pairs with less than 15 nm spacing between the projected positions of LSU and body. We estimated the density of relative orientations (ROs) in rotation-vector space, where each rotation is represented by a vector along the rotation axis with the angle as its length. Note that the rotation vector for two concatenated rotations is not, in general, given by a sum of their vectors, but for small rotations the sum is often a reasonable approximation. A raw density estimate, given by the number of RO’s in each of 512^3^ cubic voxels (0.7° on a side) was smoothed by a Gaussian (**σ** = 1.02°). The identity rotation (0° angle; central pixel) corresponds to the RO of the structural model (Natchiar et al. 2017). We used a threshold of 1.5 times the maximum density for (an equal number of) random ROs to define a region, dilated it by 2° and used it as a density-thresholding mask. Note that this is a RO mask and has to be centered on each LSU’s absolute orientation.

The 359 of 10,451 LSU/body pairs that survived RO-density thresholding were then used to create an additional constraint that uses spatial information to find bound partners as follows. While the 3D shift vector connecting the projected centers of two proteins bound in a specific conformation is characteristic for that conformation, its projection onto the image plane (the “projected shift vector”) varies in length and direction with the absolute orientation of the bound complex. We used the fact that, if complex orientation and 3D shift vector are both known, the projected shift vector can be precisely predicted and, by comparing it to the observed vector, exploited to reject FP pairs. We determined the 3D shift-vector’s constant part and also its dependence on the RO (in 1st order) by minimizing the mean-squared prediction error for the projected (2D) vector across all pairs that survived density thresholding. The resulting predictor was then used to filter candidate pairs. As upper limits for the discrepancy we used 0.35 nm for LSU/body.

A similar procedure to restrict the search space was used for body/head pairs. We again used RO density thresholding to select pairs (84/3,425) and estimated the shift-vector predictor (different from the one for LSU/body pairs), but used as a search mask a cylinder (boundaries shown in Fig. 3C) with a long axis axis matched in direction to the first principal component of those 84 pairs and a radius (5°) chosen ad hoc when inspecting the data. The cylinder’s length (32°) was extended beyond the largest angles seen among those 84 pairs to provide a control volume along the main swivel direction. The stringency of the shift-vector filter was set to 0.40 nm for body/head pairs.

The data for Fig. 3C were obtained by performing a search for heads in all 87 FOVs using both the cylinder to confine orientations and the shift-vector discrepancy as constraints. The distinction between 80S and 40S particles used results from LSU/body pair searches. The data shown in Fig. 3D and later were obtained by first searching the surround of each LSU found in any FOV for a bound body. In the next step the surround of each LSU-bound body was then searched for a head with a consistent RO (see Supplemental Figure 3). Both searches took advantage of rotation as well as shift vector constraints. Constrained searches were performed using orientions on an equally spaced (0.7° for LSU/body and 1.1) grid in rotation-vector space.

Spatial distributions of ribosomes *in situ* (Figs. 1C and 3E) were rendered in PyMol (v. 2.2, Schroedinger). To place full ribosomes (LSU, body, and head) in renderings, the reference model including all three components (PDB: 6ek0) was rotated and shifted to bring the LSU component into agreement with the cHRTM-derived LSU orientation and image-plane location (x,y), and then shifted along the axis perpendicular to the image plane (z) by an amount equal to the optimized template defocus. The body and head components of the model were then additionally rotated and shifted to bring each into agreement with the cHRTM-derived orientation and image-plane (x,y) locations. A similar approach was followed to place free SSUs (not near any LSUs; Fig. 1C).

### Ligand detection

We generated a set of search templates that were not only specific for particular ligands but also were matched individually to the known LSU orientations and locations for each instance, such that the peak CC signal will appear at a particular pixel in the CCG. To create a reference structure for a targeted ligand we aligned the model that contained that target (including both 80S ribosome and any bound ligands) to our core LSU model. Rigid-body rotations and translations were performed using Pymol (v. 2.2, Schroedinger; 5 refinement cycles, outlier rejection cutoff set to 2.0 A; alignment mode=“super”), and the ligand was then extracted from the aligned structure. cHRTM searches limited to a single template were performed on the image patches surrounding the locations of detected ribosomes; because only one orientation was tested, CCGs were first calculated for 1000 random orientations to estimate the mean and SD required for flat-fielding. When needed (Fig. 4D), the signals from multiple instances were combined by averaging the CCGs. To estimate the combined value’s uncertainty we used the fact that the flat-fielded signals have a standard deviation of one and made the assumption of independence. We estimated an occupancy value and its SD by dividing both by the SNR expected for the target’s MW and the local sample thickness. The 80S models with bound ligands that we tested here (see Supplemental Figure 4) were: A/T-site tRNA, Protein Data Bank (PDB) code: 4uje (Budkevich et al. 2014); A/A-site tRNA, 6om7 (Li et al. 2019); A/P-site tRNA, 6om0 (Li et al. 2019); P/P-site tRNA, 6olz (Li et al. 2019), P/E-site tRNA, 6om0 (Li et al. 2019); E/E-site tRNA, 6r5q (Shanmuganathan et al. 2019); A/P*-site tRNA (chimeric), 6gz3 (Flis et al. 2018); P/E*-site tRNA (chimeric), 6gz3 (Flis et al. 2018); elongation factor 1A, 4uje (Budkevich et al. 2014); elongation factor 2, 6gz3 (Flis et al. 2018).

The regions shown in Figure 4 are a Voronoi parcellation of the intersubunit rotation (ISR) *vs*. head swivel plane that was used to group ribosomes for ligand detection and to compare conformational prevalence for different conditions. These regions were generated using a set of seed points based on the unrotated model for region 1; ROs determined between our reference structure and each of three published reconstructions for regions 2-4 (Behrmann et al. 2015), and two atomic models for regions 7 and 8 (Flis et al. 2018). Region 6 was defined using the coordinates at the midpoint between two atomic models ((Brown et al. 2018), (Voorhees et al. 2014)). Where no structures were available, equally spaced points between the nearest defined regions were used (region 5,9,10).

### Estimating jdMW

Attenuation constants for scattering losses were determined as described by (Feja and Aebi 1999). Scattering lengths that we found were 283 nm for total loss (with a 100 μm aperture) and 314 nm (no aperture (Rice et al. 2018); see Supplemental Figure 2). The attenuation in SNR with sample thickness, T (Fig. 2D), was determined by fitting a set of exponentials with amplitudes left free but allowing only one decay constant, λ_SNR_, for all fragment SNRs. The jdMW was then estimated by fitting the following function: jdMW = ((SNR_thr_*exp(T/λ_SNR_)-b) / A)^2 to determine values for the free parameters A and b. For a typical search, where we assume that an SNR_thr_ of 8 is required to achieve an acceptable false-positive rate after optimization, A=0.47 and b=2.09 provided a good fit.

## Author contributions

J.P.R. and W.D. designed the study, with essential input from J.L.-S, and wrote the paper. J.P.R. performed all experiments with essential help from H.C. and analyzed all data with input from W.D.

## Acknowledgements

We are grateful for the help from Zhiheng Yu and the Janelia Cryo-EM facility, and from Christian Guggenberger from the MPG’s Garching computing facility. We wish to thank Niko Grigorieff, Tim Grant and Ben Himes for discussions during the early part of the project, Peter Detwiler, Tim Ryan, Ilme Schlichting, and Rob Singer for helpful comments on the manuscript, and Julia Kuhl for help with graphics.

## Data and materials availability

A dataset including all intracellular ribosome images contributing here is in preparation and will be made publicly available at EMPIAR (https://www.ebi.ac.uk/pdbe/emdb/empiar/).

## Supplemental Figures S1-S5

**Fig. S1.**
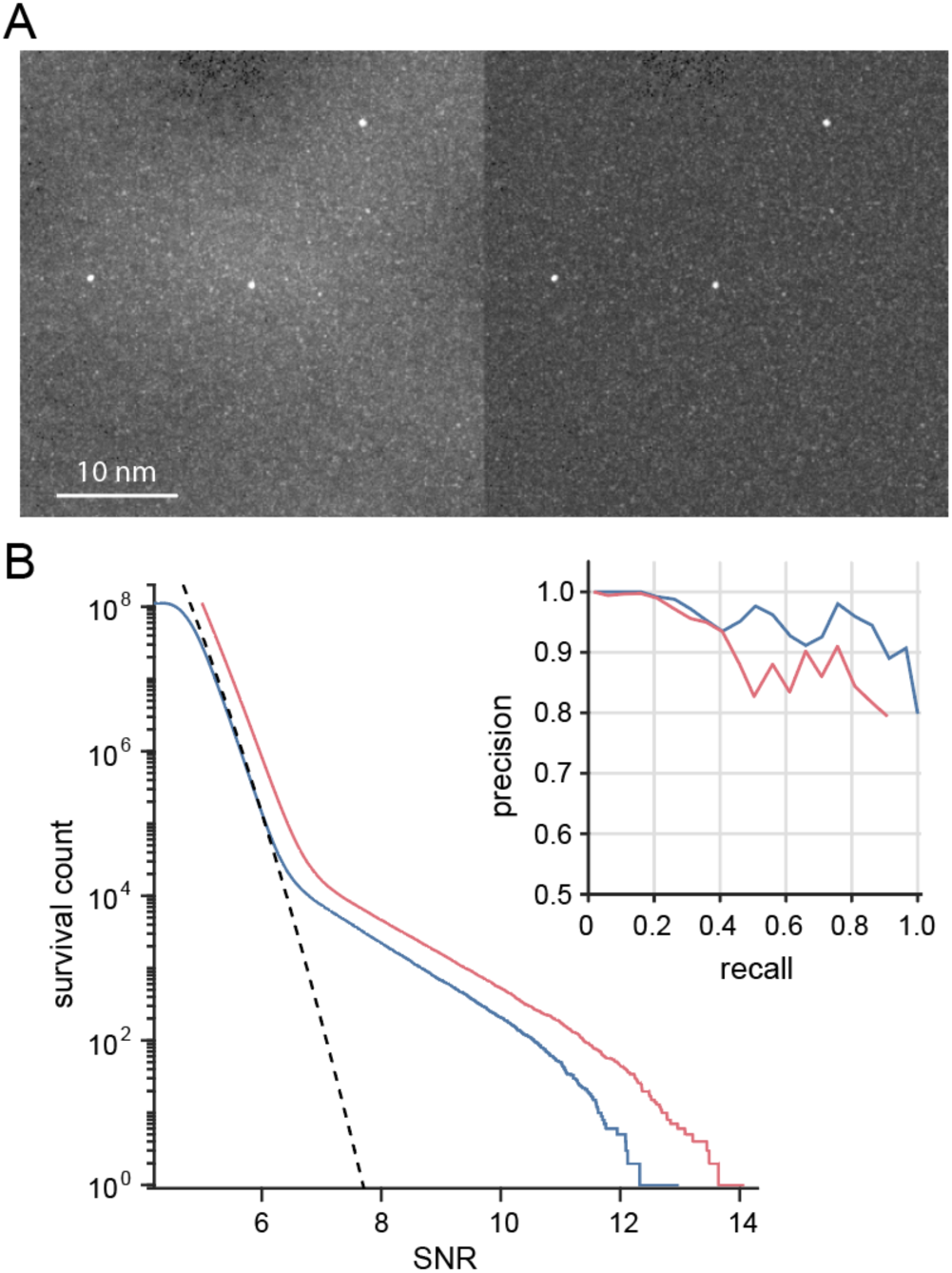
Flat-fielding. **A)** Maximum intensity projection of CC values across orientation and sample depth from a search for the LSU before (left image) and after (right image) flat-fielding. **B)** Amplitude distributions as survival histograms before (red) and after (blue) flat-fielding. Also shown: Gaussian noise (dashed). Sample thickness within this image varied between 195-295 nm (first and last deciles). Inset: precision *vs*. recall before (red) and after (blue) flat fielding for 1145 instances in 24 images with a sample thickness range from 50 to 320 nm (mean ≈ 150 nm).

**Fig. S2.**
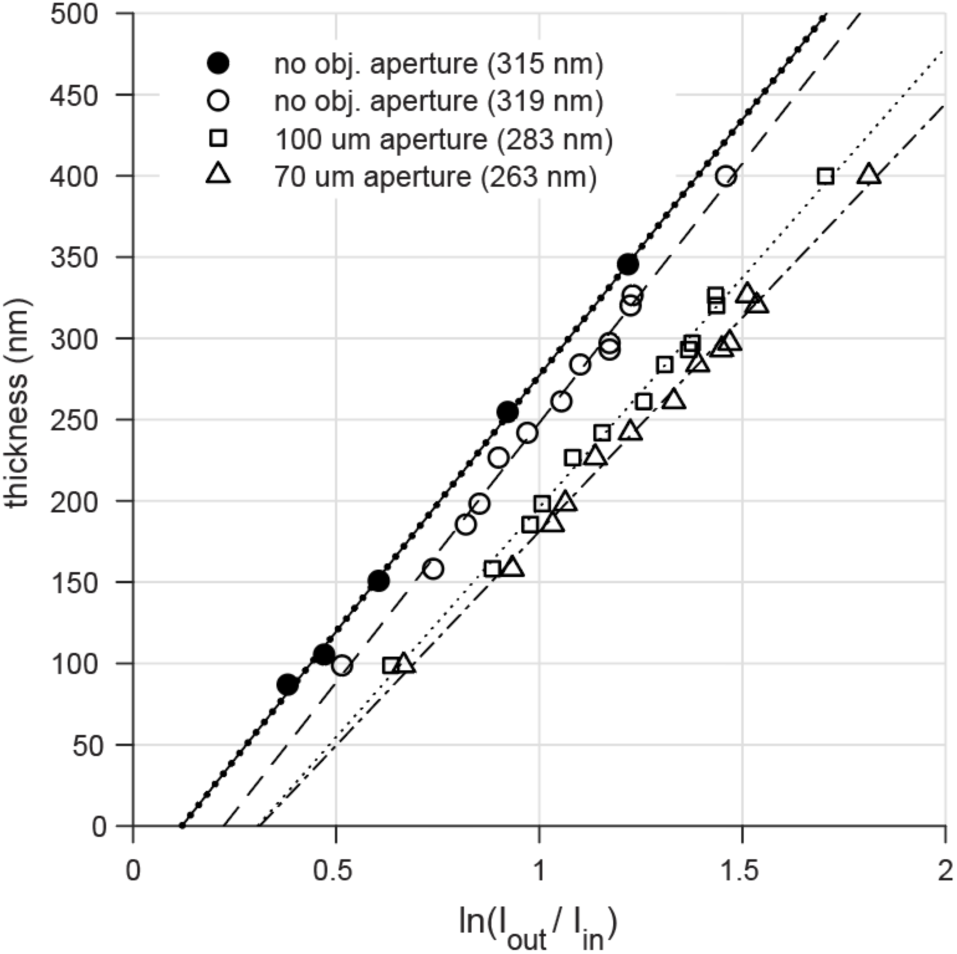
Calibration of the relation between sample thickness and relative image brightness with the energy filter slit removed (I_out_) and inserted (I_in_); filled symbols: Krios 1, MEF cells; open symbols: Krios 2, U2OS cells. Fitted slopes in parentheses. For details see Ref. (Feja and Aebi 1999).

**Fig. S3.**
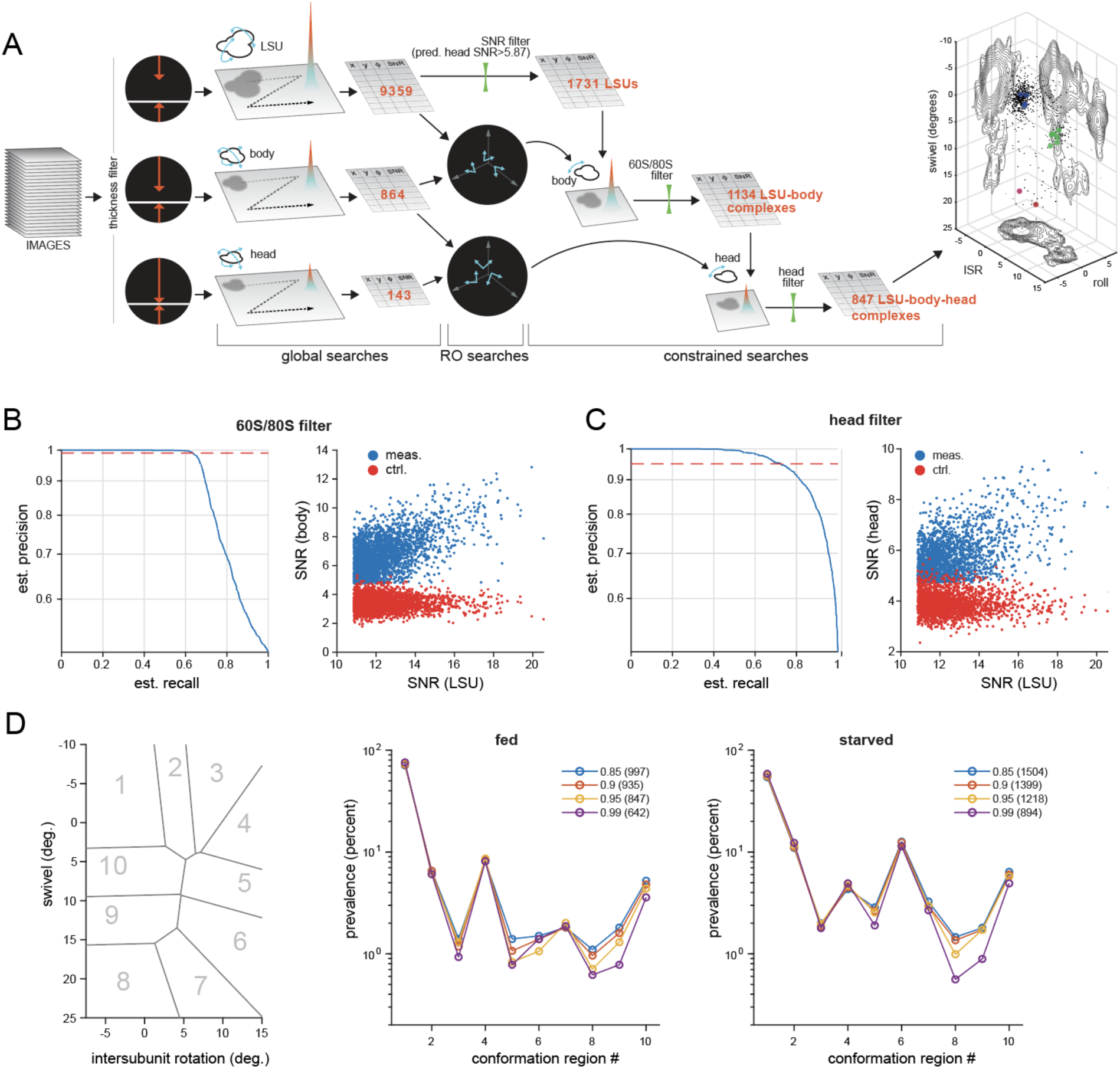
**A)** Whole ribosome search, schematic. From the left: set of motion-corrected images, sample-thickness thresholding, unrestricted search, relative orientation cluster detection, focused search for bodies using LSU position and orientation information (center lane). Bound/unbound selection. Focussed search for heads, output. **B-C)** Precision *vs*. recall for LSU-bound detection of bodies and heads, created by varying the SNR threshold to filter results from constrained searches until the estimated detection precision dropped below thresholds of 0.95 and 0.99, respectively. Detection precision was estimated for each SNR threshold as the number of detected instances in the target’s search range (the signal channel; presumed TPs), divided by the sum of TPs and presumed FPs found in control searches (the noise channel). **D)** Prevalence *vs*. conformation (regions are indicated at left), shown for several different estimated detection precisions. Legend: estimated precision; number of 80S ribosomes detected. Shown for normally-maintained and nutrient-starved cells.

**Fig. S4.**
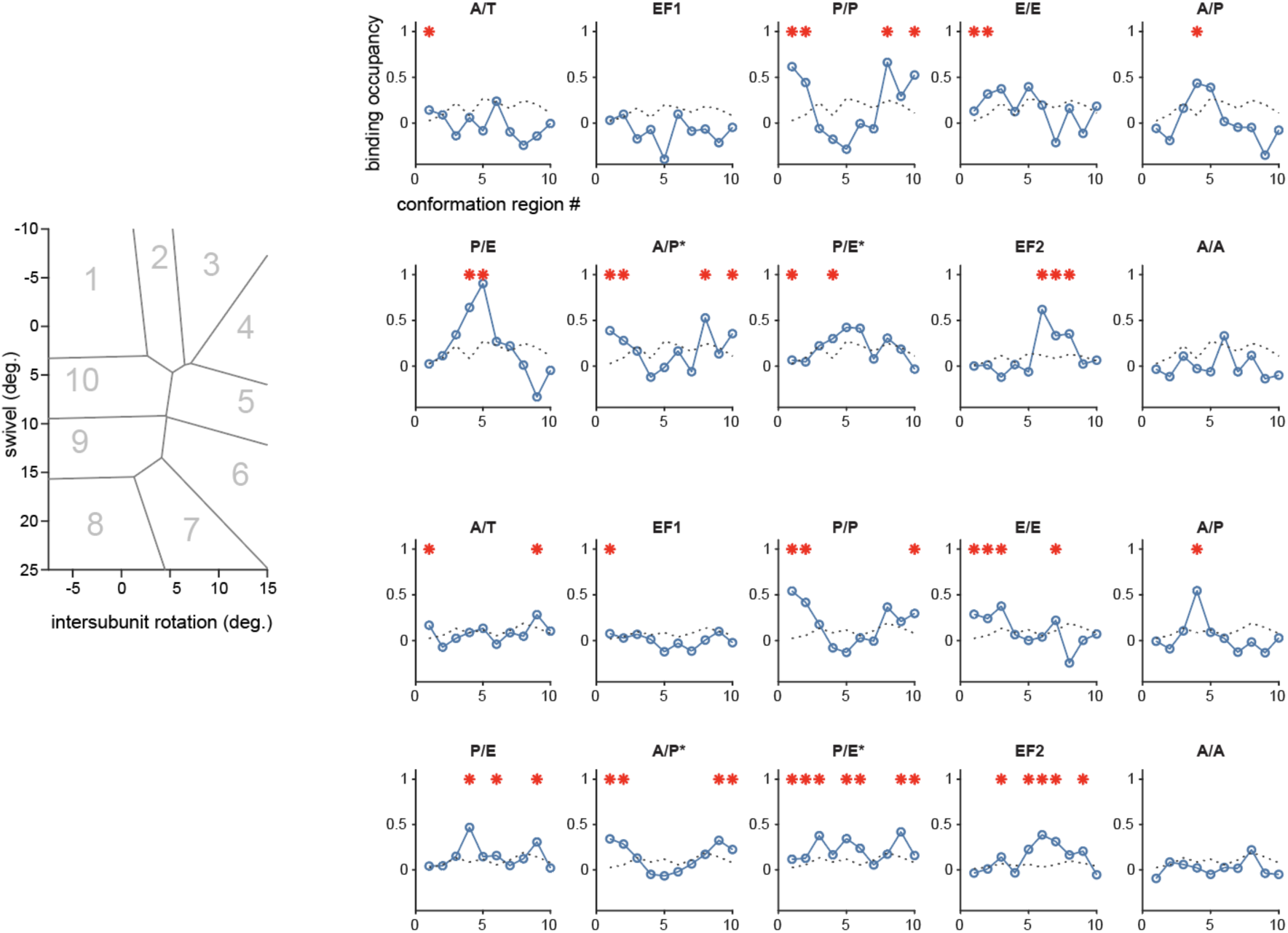
Estimated ligand-binding occupancies for 80S ribosomes in the conformations indicated (left), shown for normally-maintained (upper) and nutrient-starved (lower) cells. Dotted lines show the standard deviation (S.D.) in each region; red asterisk indicates occupancy > 2 S.D.

**Fig. S5.**
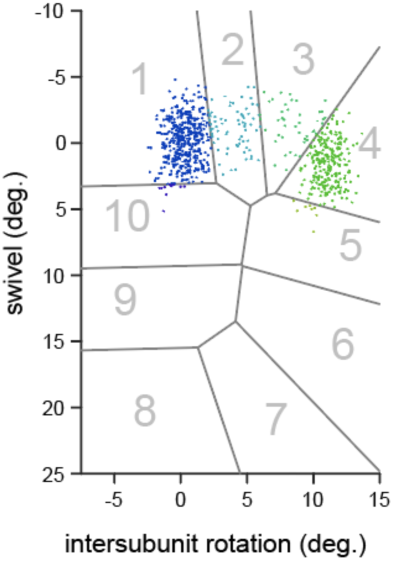
Points generated from the conformation parameters described in Ref. (Behrmann et al. 2015) by adding Gaussian noise (**σ** = 0.9 degrees, 1.8 degrees for intersubunit rotation and swivel, respectively) matched in spread to what we see in region 1 of our dataset, which is probably caused by a combination of measurement noise and actual conformational fluctuations. The total number of points for each conformation reflects the prevalence given to it in Ref. (Behrmann et al. 2015).

